# Associative transcriptomics in *Brassica napus* suggests a role for Arabidopsis Response Regulator orthologs in seedling vigour

**DOI:** 10.64898/2025.12.03.692010

**Authors:** Laura Siles, Guillaume N. Menard, Rachel Wells, Ange Zoclanclounon, Peter J. Eastmond, Smita Kurup

## Abstract

**Background and Aims:** Optimal seedling vigour determines successful crop establishment and its ability to overcome abiotic and biotic threats. Seedling vigour is regulated by several different factors, genetic, physiological, environmental, post-harvest storage, making it challenging to fully elucidate. The aim of this study was to dissect the genetic basis of post-germinative seedling traits and its implications in seedling establishment success in *Brassica napus*.

**Methods:** A *B. napus* diversity set panel was used to phenotype seedling establishment traits in the soil including days to emergence, days to appearance of first and second true leaves, and days to reach 25% and 50% cotyledon expansion. An associative transcriptomics analysis was performed to identify the genetic factors regulating seedling establishment.

**Key Results:** The study of different phenotypic traits highlighted the relevance of scoring more than one trait to determine good performers. Gene expression markers significantly associated with the studied traits were associated with the cytokinin signalling pathway and photomorphogenesis. Gene expression markers for orthologues of *Arabidopsis thaliana RESPONSE REGULATOR 4* and *5 (ARR4, ARR5)* were significantly associated with days to appearance of second leaf. *SUPPRESSOR OF PHYA-105 1* (*SPA1)* orthologues were associated with cotyledon area, meanwhile *ALTERED SEED GERMINATION 7* (*ASG7*) and *PSEUDO-RESPONSE REGULATOR 7* (*PRR7)* orthologues were associated with days to emergence, suggesting a complex and interconnected regulatory pathway controlling several traits of seedling establishment.

**Conclusions:** These results uncovered promising genes that appear to work in a complex and interconnected manner controlling several post-germinative seedling traits affecting final seedling establishment speed.

## Introduction

The first critical step for good crop production is successful seedling establishment. Good seed/seedling vigour (henceforth referred to as seedling vigour) is defined as the sum of seed properties that determine the ability of seeds to germinate and establish rapidly and uniformly (Redona and Mackill, 1996; Finch-Savage and Bassel, 2015; Basu and Groot, 2023). Due to its agronomic importance, seedling vigour has been investigated in many crop species. Studies in rice, wheat, maize and Brassica species have focused on seed germination and dormancy (Hrstková et al., 2006; Wang et al., 2010; Hatzig et al., 2015; Gu et al., 2017; Zhao et al., 2021), as well as seedling length in terms of shoot and root length (Finch-Savage et al., 2010; Basnet et al., 2015; Lee et al., 2017; Zhang et al., 2017; Menard et al., 2021; Zeng et al., 2022). Seedling vigour is determined by environmental and genetic components. While environmental and maternal effects have been extensively studied (Awan et al., 2018; Chen et al., 2021; Sharma et al., 2022; Bailly and Gomez Roldan, 2023; Tarnawa Á et al., 2023), the genetic factors influencing this trait are less well characterised because of the different number of factors in play. Specifically, the genetic determinants and variations for seedling vigour and biomass accumulation in Brassica crops are still little understood (Basnet et al., 2015; Nelson et al., 2022).

Despite oilseed rape (*Brassica napus*, L., OSR) being the third largest source of vegetable oil in the world and a crucial source of biofuel production (Food and Agriculture Organization of the United Nations, 2021), its yield has not increased substantially since 1980. Enhanced seedling vigour and establishment are desirable traits for all OSR varieties; traits that can help increase its final crop yield. In addition, well established seedlings have a higher opportunity of survival and lower vulnerability to factors such as low precipitation, droughts, weeds and plant diseases (Kutcher et al., 2010; Assefa et al., 2018; Ortega-Ramos et al., 2021). Rapid seedling development and the subsequent appearance of the first true two and four leaves are key traits to overcome pest attacks, particularly cabbage stem flea beetle, the most significant biotic threat to OSR cultivation in Europe (Jordan et al., 2020; Zheng et al., 2020; Ortega-Ramos et al., 2021). An earlier onset of autotrophic growth, increased levels of the phytohormones auxin, cytokinins, brassinosteroids and gibberellins have been linked to an increase in seedling vigour in *Arabidopsis thaliana,* as well as OSR (Spartz et al., 2012; Sahni et al., 2016; Boniecka et al., 2019; Boter et al., 2019; Zhu et al., 2020; Sharma et al., 2022).

To uncover the genetic bases of trait variation, different methods including Associative Transcriptomics (AT), can be applied to uncover genes controlling these traits. AT is a robust method successfully used in crops that utilises RNAseq to associate both sequence variants and gene expression levels with phenotypic trait data (Harper et al., 2012; Miller et al., 2016; Havlickova et al., 2018; Woodhouse et al., 2021; Sahu et al., 2023). Here, we used AT analysis to identify genes involved in seedling vigour in a diversity OSR set panel that included a wide range of crop types. Seed germination, time to emergence, cotyledon expansion, days to appearance of first and second leaves as well as hypocotyl and root length were recorded. The main objective of this study was to obtain a better understanding of the genetic control of OSR seedling establishment, specifically days to appearance of second true leaf, and the relationship of this key trait with other seedling vigour traits to improve seed establishment speed and uniformity.

## Material and Methods

### Plant material

A *B. napus* diversity subset of 95 genotypes from the BnaASSYST panel was used to provide a representation of the breadth of this species diversity (Harper et al., 2012; Havlickova et al., 2018). All seeds were obtained from plants grown at the same time under glasshouse conditions to avoid environmental effects. Individual genotypes were classified as Winter OSR, Spring OSR, Semiwinter OSR and Others (which included swede, kale, unspecified and fodder genotypes) [**Supplementary information Table 1**]. Eight pre-breeding lines were also included in the study designated Line 1-8. These lines were provided by a breeding company (see Acknowledgements) and were known to exhibit good seedling establishment; hence they were included as a control subset. In all, a total of 103 genotypes were scored in the experiment.

**Table 1.**
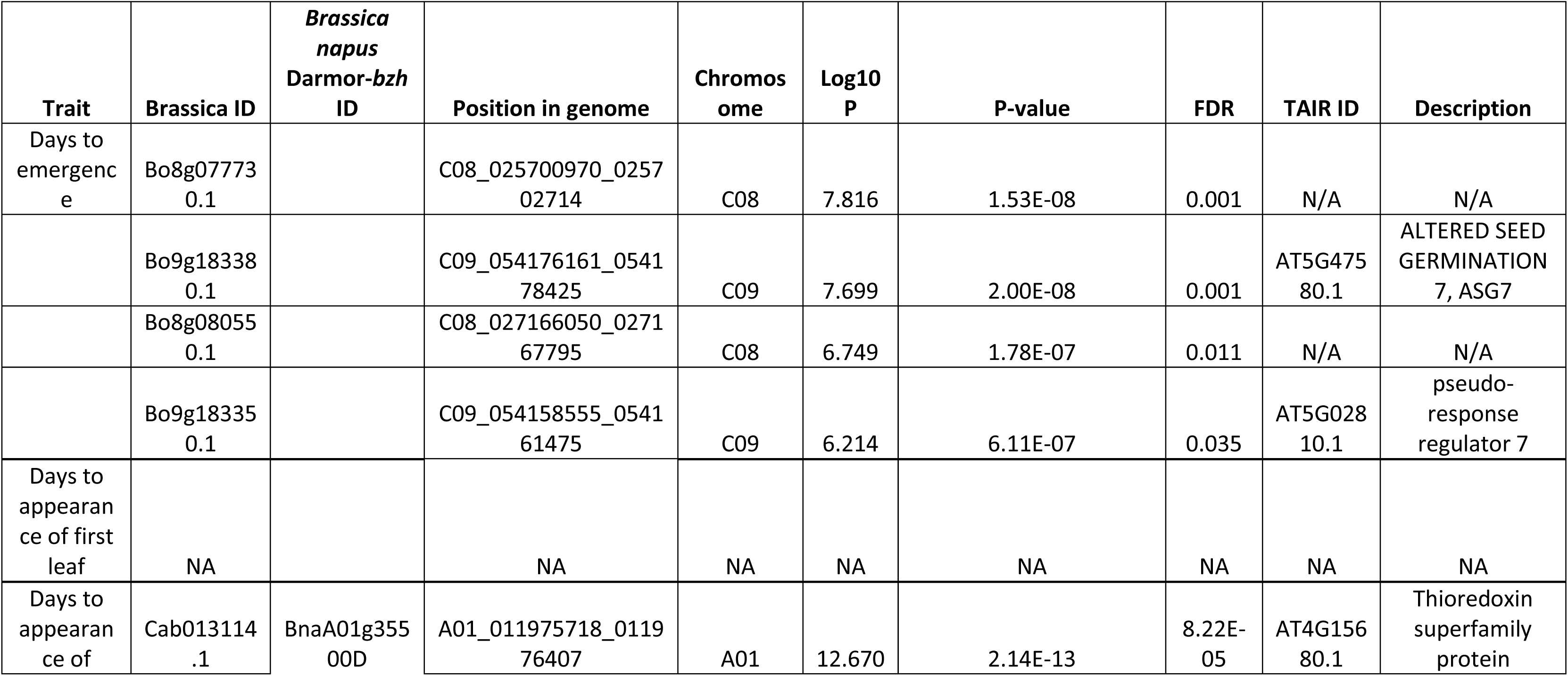
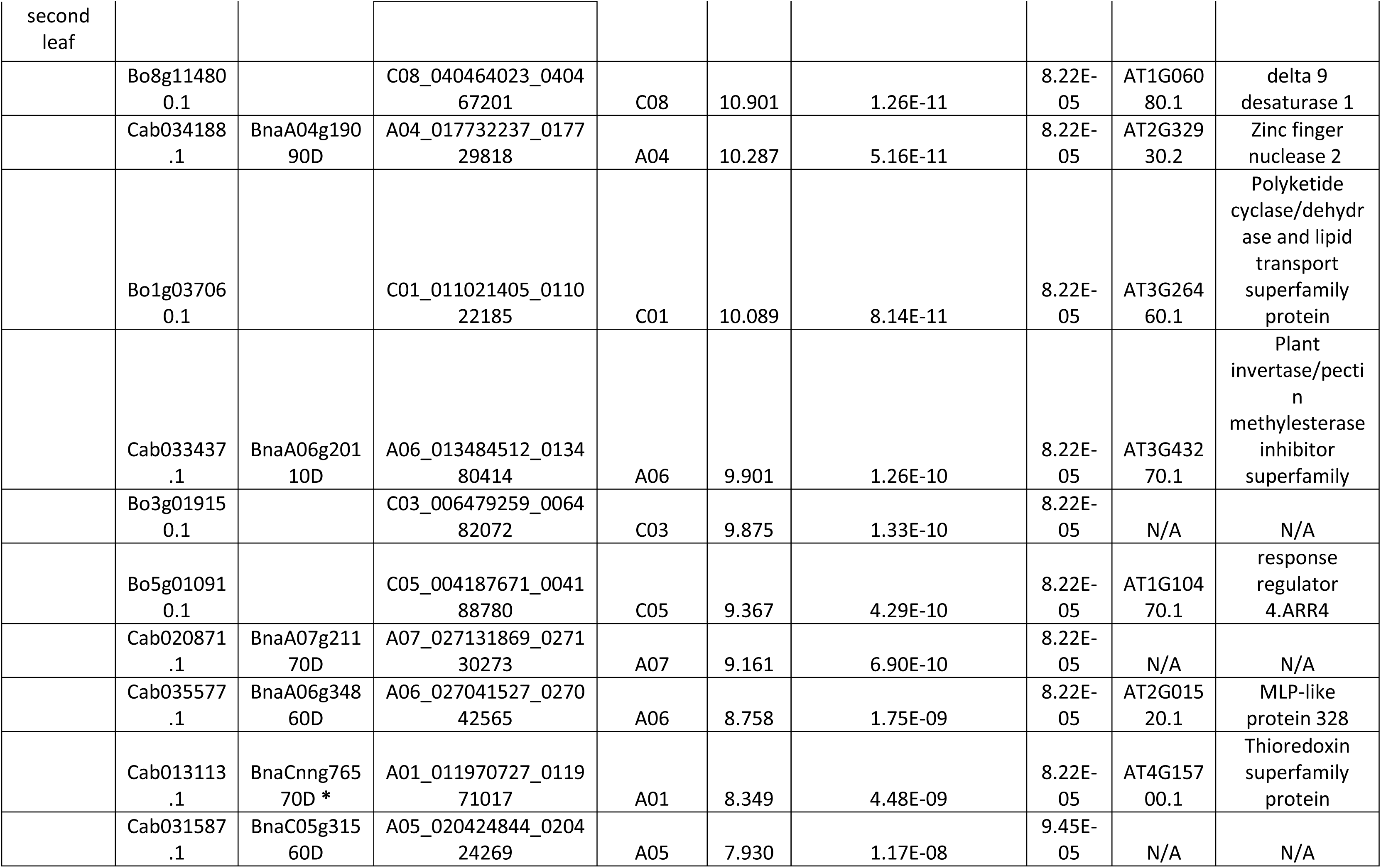
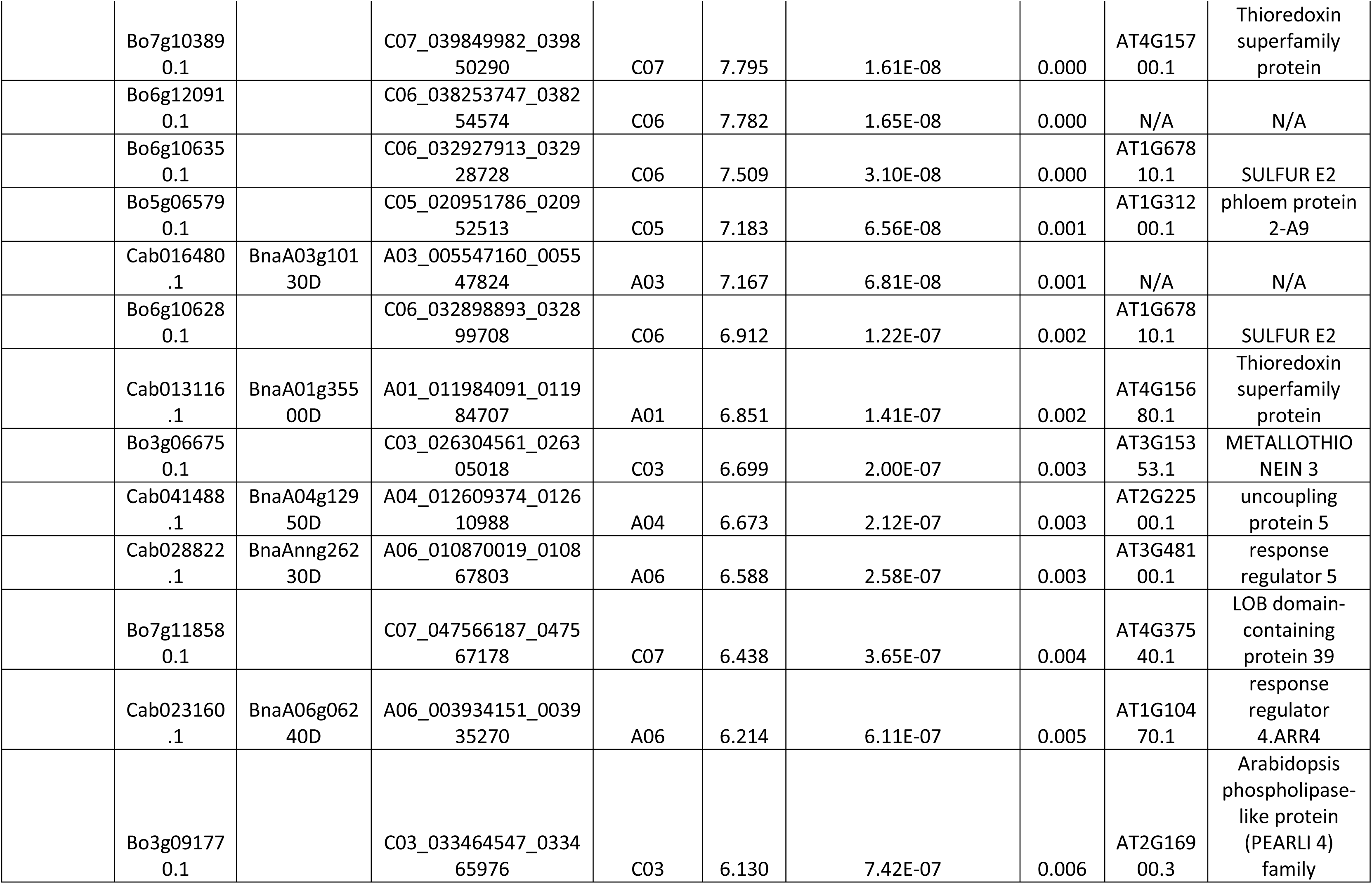
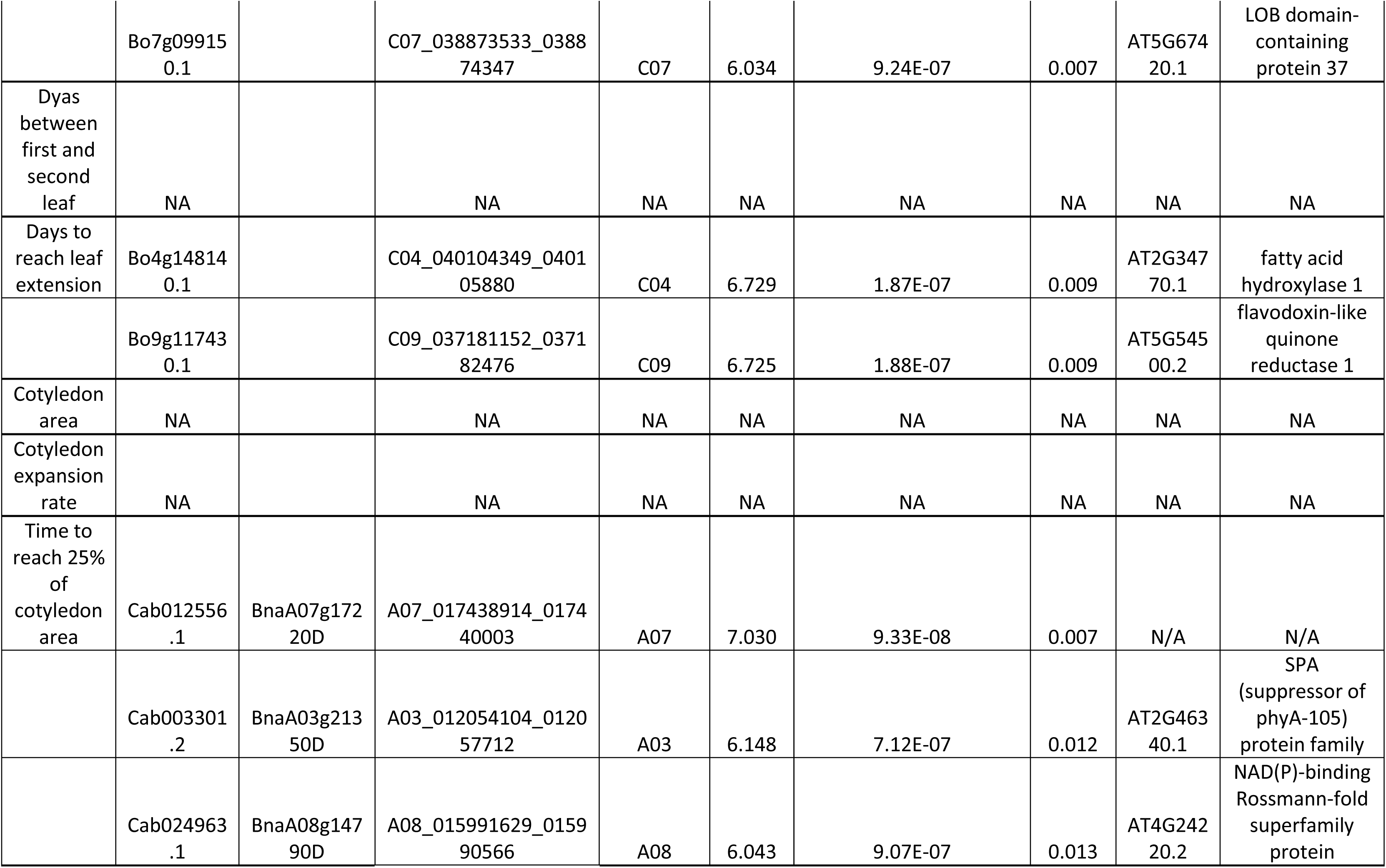
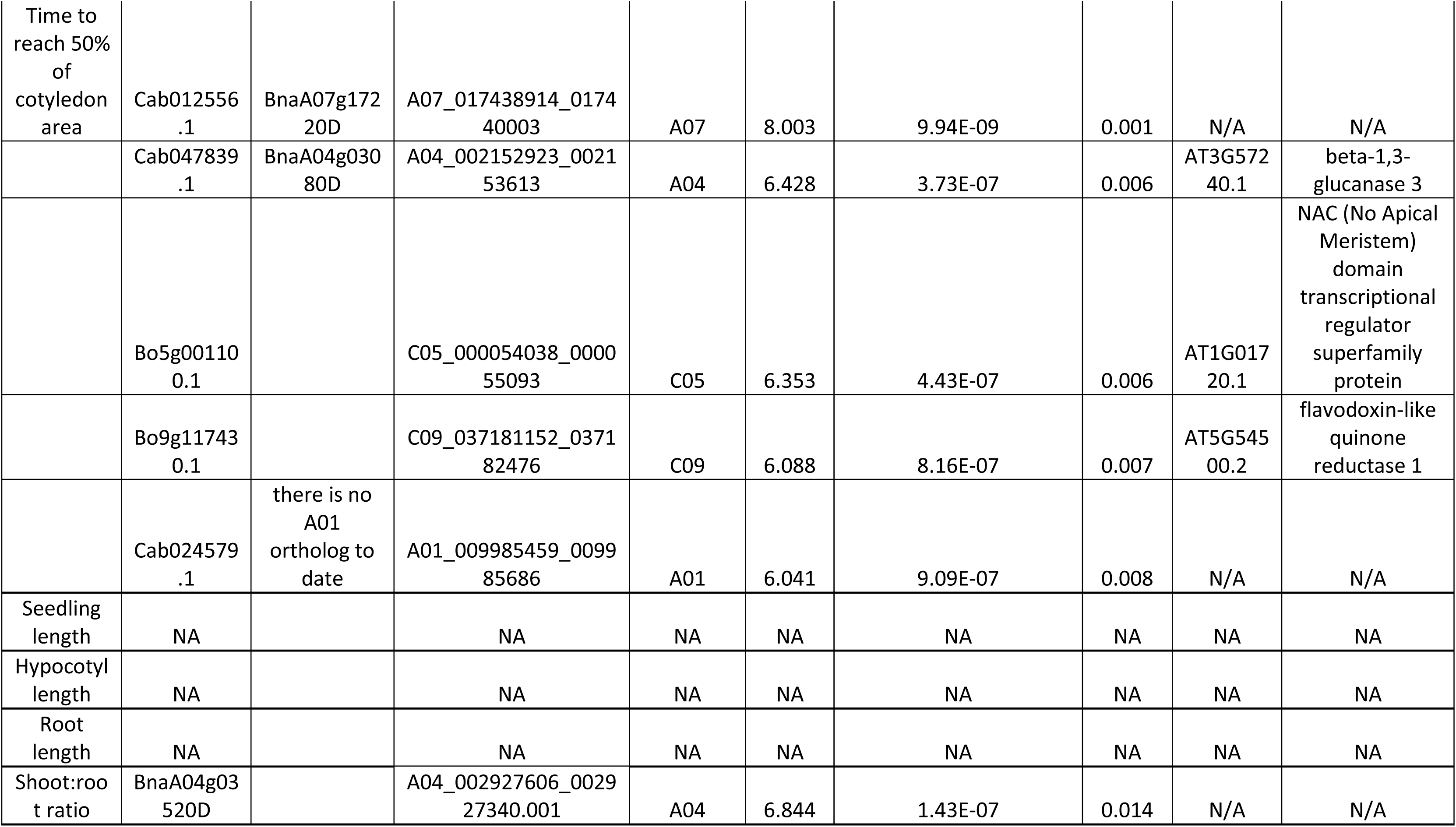
Gene expression markers (GEMs) associated with measured seedling traits over Bonferroni threshold (6.03). Brassica ID corresponds to Pan-transcriptome gene ID http://yorknowledgebase.info. NA represents no significant GEM association with the measured seedling trait. N/A was added when no Arabidopsis homolog was found. *Brassica napus* cultivar Darmor-*bzh* ID is shown from blast results. \**Brassica napus* orthologs from Ensembl Plants. Position of each gene in the genome comes from Havlickova et al., 2017 (https://doi.org/10.1111/tpj.13767), Appendix S2

### Seedling growth in media

OSR seeds were placed on a 25 cm square polypropylene culture plate containing sterile Gellan Gum gel (FUJIFILM Wako Pure Chemical Corporation) at 2 g/L in ¼ x Murashige and Skoog media (pH 5.0, Sigma-Aldrich, Missouri, United States) in deionized water. Calcium chloride at a final concentration of 4 mM was added to solidify the media previous to pouring 250 ml of media in each plate. A defined grid was used to ensure consistent positioning of the seed across the plate, between plates and replicates. Each plate contained 2 genotypes, with 10 seeds per genotype and a total of 20 seeds per plate. Plates were sealed using micropore tape and placed in a sealed light tight black bag (BLK1215HEAVY 300 mm × 375 mm × 100 micron, Polybags UK) to ensure complete darkness. Plates in batches of 32 were arranged upright (80°) on plastic supports [**Supplementary Information Figure 1A**]. Plates were kept in complete darkness for 3 days at constant 20°C and 60% relative humidity (SANYO growth cabinet). After 3 days in darkness, the black bags were removed and the plates were kept upright and arranged in a randomised statistical design on the plastic support. The growth conditions were adjusted to 16 hours day, 8 hours night, 100 µmols m^−2^ s^−1^ of light intensity and constant temperature of 20°C for a further 2 days. After 5 days, the plates were photographed using a high-resolution camera (NIKON D5300 with NIKKOR 50mm f1.8 prime lens, Minato, Tokyo, Japan) mounted on a photobench equipped with additional lights. Seedling traits were measured using the SmartRoot tool in FIJI (Schindelin et al., 2012). Root length was measured from the collet (root–hypocotyl junction) to the root tip, meanwhile hypocotyl length was measured from the collet to the shoot apical meristem. The seedling length was the sum of hypocotyl and root length. The 40 blocks were divided in 5 runs each of 8 blocks. All 103 genotypes were replicated 3 times, with a total of 30 seeds per genotype. Data was transformed to ensure homogeneity of variance when necessary. The data was analysed as a linear mixed model using the REML algorithm (asreml-R v3.0 and Genstat 22^th^ Edition), with the fixed effect of genotype assessed using either a Wald test (assessed against a chi-square distribution) or a variance ratio test (assessed against an F distribution).

**Figure 1.**
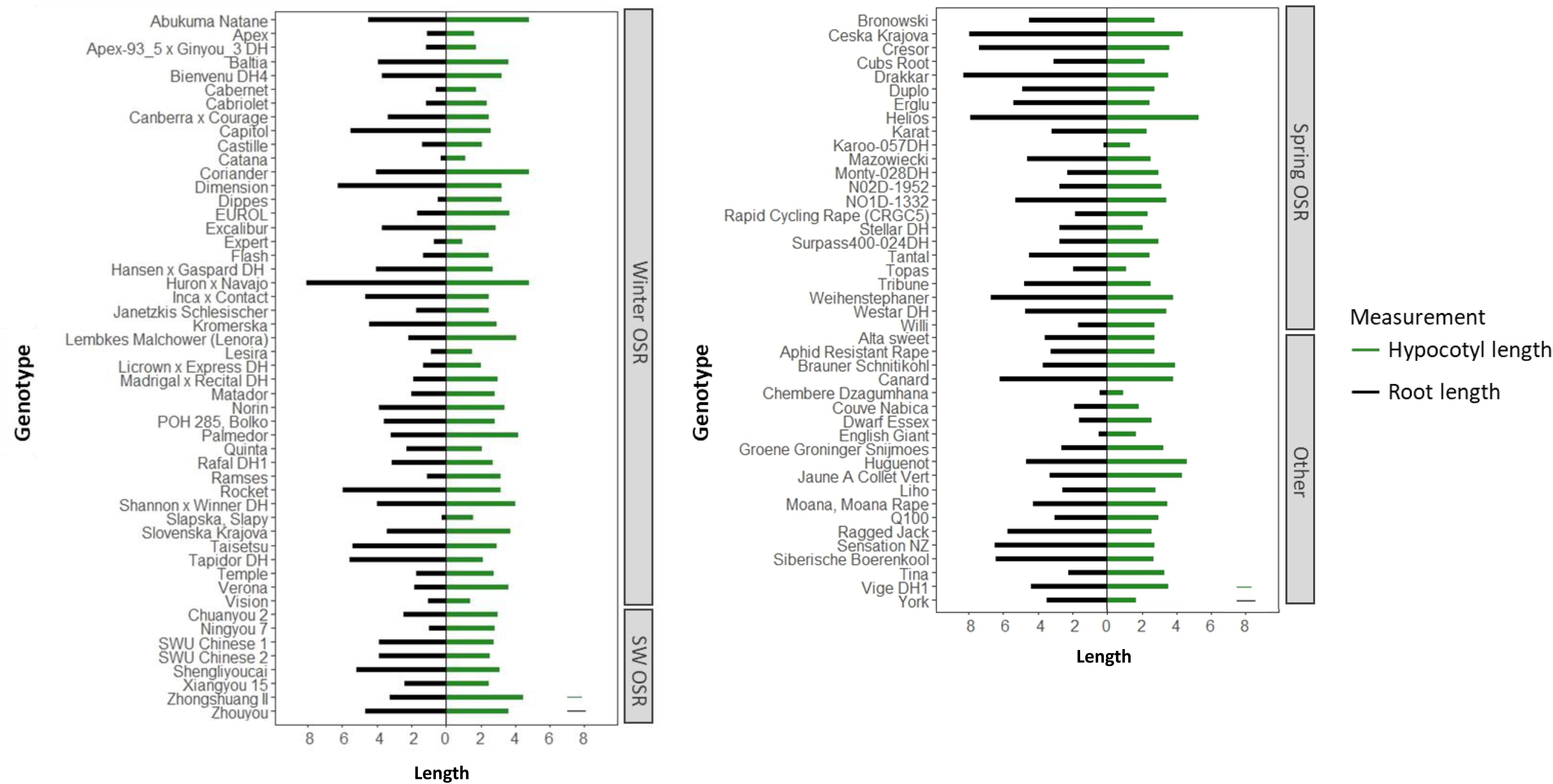
Root (black) and hypocotyl length (green) for the 103 genotypes studied divided in 5 OSR groups (Winter OSR, Spring OSR, Semiwinter OSR (SW OSR), Other and in-house lines). Data are the mean of three biological replicates with 10 seeds per replicate. Average least significant difference are represented as green and black lines in the bottom right corner of the graph for hypocotyl and root length, respectively.

### Seed germination

A total of 20 seeds per genotype were placed in each row, with a total of 15 rows per plate. Each row corresponded to a different genotype. After 3 days in complete darkness, the plates were removed from the black bags and germination percentage was recorded [**Supplementary information Figure 1B**]. Seeds were considered to have germinated when the radicle was 3 mm long. All 103 genotypes were replicated 3 times in a resolvable latinized row-column alpha design with 3 replicates and 7 blocks per replicate with 15 positions per block, ensuring that each genotype occurred at most once at any of the 15 positions across the whole experiment. The data was analysed as a linear mixed model using the REML algorithm, with the fixed effect of genotype assessed using either a Wald test (assessed against a chi-square distribution) or a variance ratio test (assessed against an F distribution).

### Seedling vigour in soil

One seed per pot was sown at a depth of 1 cm in a 0.35 litre pot containing Rothamsted Prescription Mix: 75% medium grade peat, 12% screened sterilized loam, 3% medium grade vermiculite, 10% grit (5 mm screened, lime free), 3.5 kg m−3−3 Osmocote Exact (total N 16%) (supplier: Scotts UK Professional, Ipswich, Suffolk, UK) and 0.5 kg m−3−3 PG compound fertilizer (supplier: Yara UK Ltd., Harvest House, Europarc, Grimsby, N E Lincolnshire, UK). A total of 14 pots were placed in each tray, all 15 trays were placed in a SANYO growth cabinet with 16 hours light, 8 hours dark, 16°C day, 15°C night, 60% relative humidity and 190 µmol m^−2^ s^−1^ of light intensity. Each genotype had 4 biological replicates. Days to emergence was scored when the seedling hook could be observed emerging from the soil. ‘Days to appearance of first leaf (DAFL)’ was measured from emergence until the first true leaf could be visually observed emerging from the main meristem. ‘Days to appearance of second leaf (DASL)’ was measured from emergence until the second true leaf could also be visually observed emerging from the main meristem. Seedling growth was followed until the first true leaf reached 3 cm long and the second true leaf reached 2 cm long. To measure cotyledon expansion and maximum cotyledon area, individual trays were photographed as previously described. Images were processed using FIJI. The experiment was arranged as a non-resolvable incomplete block design over two occasions, and when necessary, traits were first transformed to ensure homogeneity of variance. Each occasion had 15 trays, and each tray contained 14 pots. The data was analysed as a linear mixed model using the REML algorithm, with the fixed effect of genotype assessed using either a Wald test (assessed against a chi-square distribution) or a variance ratio test (assessed against an F distribution).

#### Seed size

One hundred mature seeds from the whole plant were placed in a petri dish and images were acquired using a Videometer (Videometer A/S, Herlev, Denmark). For obtaining seed area, three technical replicates for each plant were performed, and seed area was averaged for each plant.

### Associative transcriptomics analysis

The predicted means for all the measured seedling vigour traits were analysed and run through a Gene Expression Marker (GEM) association and SNPs pipeline R package, the GAGA, available at https://github.com/bsnichols/GAGA (https://zenodo.org/records/7034543). GEM association analysis was performed using linear marker regression using Reads per Kilobase of transcript, per Million mapped reads (RPKM). All GEMs with an average expression across the population of less than 0.5RPKM were excluded during analysis. The significance thresholds for trait association was determined using the false discover rate (FDR). Bonferroni threshold was calculated as - log10(0.05/number of markers). In this study we had 53883 markers for GEMs, with a Bonferroni threshold of 6.03. The genotype, expression level datasets and reference used (Pantranscriptome, version 11) were published by Havlickova et al. (2018) and are available at http://www.yorknowledgebase.info/. The GAGA pipeline also uses GAPIT for GWAS (Wang and Zhang, 2021), and the three models used in this study were GLM (General Linear Models), FarmCPU and Blink. Data were passed through GAGA pipeline for GWAS but no significant SNPs associated with the traits were identified that were consistent between models.

## Results

### Phenotyping of post-germinative seedling traits

We performed a comprehensive analysis of seedling traits in OSR to have a better understanding of the individual seedling components and how these relate to seedling establishment. The genotypes tested had good germination percentages, with 80 genotypes out of 103 having a seed germination percentage over 80%, and a mean of 83.5% germination across all genotypes indicating good seed quality. In this study, we focused on post-germination traits, scored on seeds germinated in soil. However, seedling, hypocotyl, and root length; three traits that are difficult to measure in soil, were obtained from seeds grown *in vitro* (on Gellan Gum gel). To mimic seed germination in soil conditions, seeds were grown in the dark for 3 days followed by 3 days of 16-hour photoperiod. Seedling length was an extremely variable trait between the 103 genotypes, with values ranging from 1.4 to 13.2 cm. Hypocotyl and root lengths were significantly different between the genotypes tested (F_102,168_=11.28, P<0.001 and F_102,167_=17.36, P<0.001, respectively), with values varying from 0.94 to 6.38 cm and 0.17 to 8.32 cm (**Figure 1**). Statistical analyses showed a significant positive correlation between root, hypocotyl, and seedling length [**Supplementary Information Figure 2**]. To assess the fastest and slowest genotypes with regard to seedling establishment, days to emergence was recorded. Most of the genotypes took 5 to 6 days to emerge from the sowing date; meanwhile the seven slowest ones took 7 to 10 days, and up to 13 days in the case of Expert (**Figure 2**). Significant statistical differences were found for ‘days to appearance of second leaf (DASL)’ between the genotypes (Wald (100 df) = 326.72, P (chi-square) < 0.001). The genotypes that were quickest to generate the second leaf took 5 to 7 ‘days to develop from emergence (DFE)’, meanwhile the slowest ones took 9 to 10 DFE, and up to 18 days in the case of BIENVENU DH4. DASL was positively correlated with ‘days to appearance of first leaf (DAFL)’ (r = 0.71), where genotypes took between 4 to 8 days to develop from DFE (F_101,258_=1.81, P<0.001, [**Supplementary Information Figure 2A**]. Despite the time the genotypes took to emergence, 25% of cotyledon maximum area was reached before DAFL, meanwhile the 50% of cotyledon maximum area was reached very close to DAFL for most of the genotypes. Days to emergence was significatively negatively correlated with root, hypocotyl, and seedling length for winter oilseed rape (WOSR) and spring oilseed rape (SOSR) and we also observed a negative correlation between days to reach leaf extension and cotyledon expansion rate, being more significant in WOSR than in SOSR [**Supplementary Information Figure 2**].

**Figure 2.**
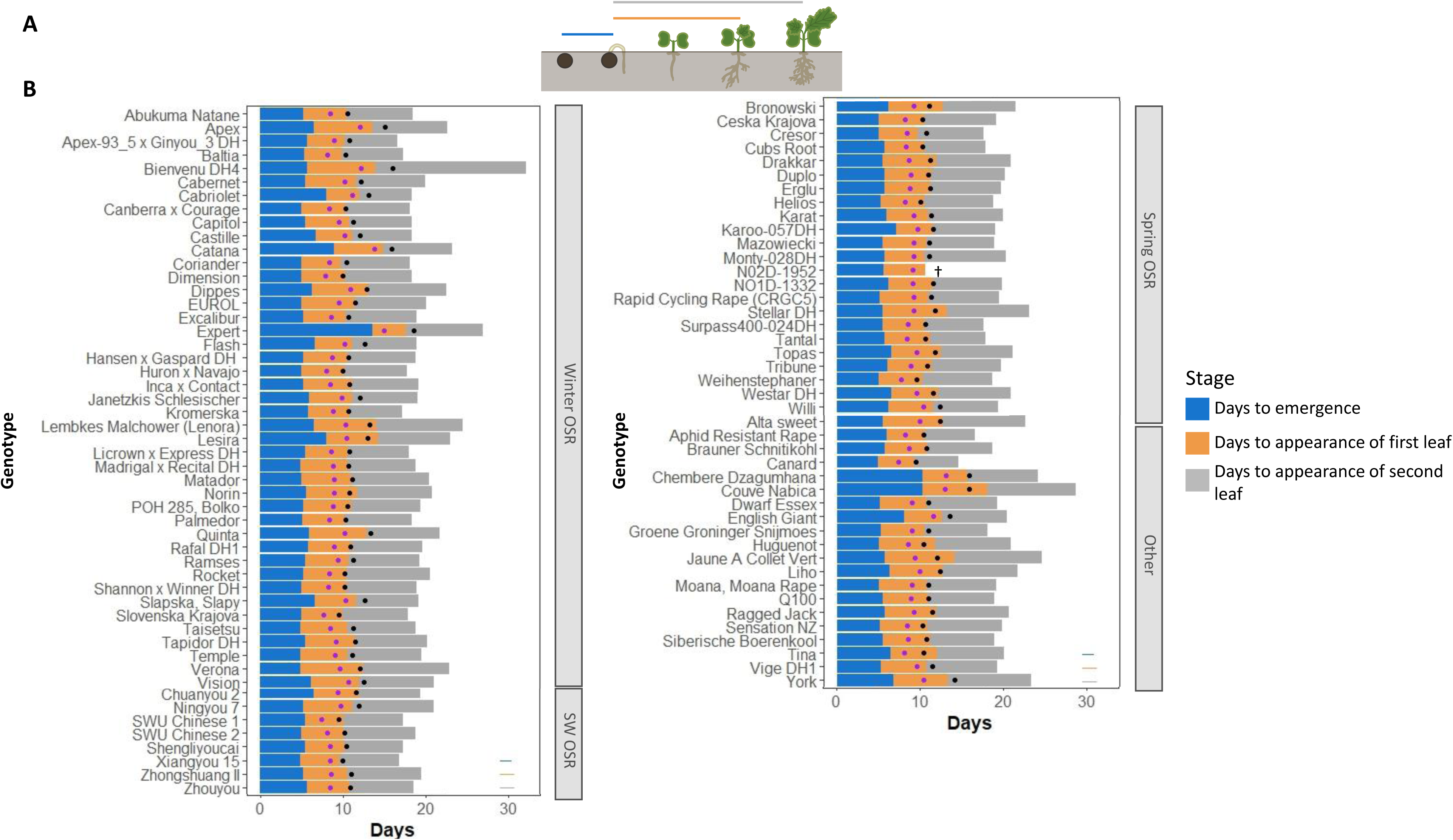
A) Graphical representation of seedling traits score: days to emergence was scored as visual appearance of hypocotyl above soil surface from sowing date. Days to appearance of first and second true leaves were scored when the first and second true leaves were visually observed emerging from the main meristem from emerging date, respectively. Bar colours represent the same developmental stage phase as the graphs in panel B. B) Days to emergence, to first and second leaf appearance for the 103 genotypes studied divided in 5 OSR groups (Winter OSR, Spring OSR, Semiwinter OSR (SW OSR), Other and in-house lines). Purple and black dots represent the day when the cotyledon reached 25% and 50% of its maximum area, respectively. Data are the mean of four biological replicates. † denotes the seedlings die before developing. Average least significant difference are represented as green, orange and grey lines in the bottom right corner of the graph for days to emergence, days of appearance of first leaf and second leaf, respectively.

### Associative Transcriptomics reveals gene expression markers associated with seedling establishment traits

To investigate the genetic bases of seedling vigour in OSR and identify significant natural variation in trait associations with gene expression levels across the population, we employed an AT analysis (Harper et al., 2012). While some phenotypic traits did not show any significant associations to gene expression levels, days to emergence was significantly positively correlated with the expression of four genes (**Table 1**). For this trait, *Bo9g183380.1* and *Bo9g183350.1* on C09 chromosome, orthologues of Arabidopsis *ALTERED SEED GERMINATION 7* (*ASG7, AT5G47580*) and *PSEUDO-RESPONSE REGULATOR 7* (*PRR7, AT5G02810*) were identified (**Figure 3B**). DASL was significantly correlated with expression of 52 genes (FDR ≤ 0.05), 25 of which exceeded the Bonferroni threshold (6.03) (**Table 1**). Candidates highly associated with this trait that appeared on more than one chromosome were identified: two orthologs of *RESPONSE REGULATOR 4* (*ARR4*, *AT1G10470*), *Cab023160.1* on chromosome A06 and *Bo5g010910.1* on chromosome C05 as well as one ortholog of *RESPONSE REGULATOR 5* (*ARR5, AT3G48100*), *Cab028822.1* on A genome (A06 chromosomes), with no copies on C genome, may be directly causally linked with DASL. These genes are involved in the cytokinin signalling pathway, and leaf expansion in Arabidopsis (Leibfried et al., 2005; Vercruyssen et al., 2014), making them promising candidates for the control of this trait (**Figure 3 C,D**). Other interesting genes whose expression is associated with this trait were *Cab013114.1* and *Cab013116.1* on A01 chromosome, two orthologs of the *MONOTHIOL GLUTAREDOXIN-S4* (*GRXS4*, *AT4G15680*). Specifically, *Cab013114.1* was the most highly associated gene expression marker (GEM) with DASL, with a -log10P value of 12.67 (Figure 3C). Therefore, these genes may be contributing to controlling the rate of appearance of the second leaf. Days to reach 25% and 50% of cotyledon maximum area also highlighted 2 promising GEMs. For 25% of cotyledon maximum area, *Cab003301.2,* an ortholog of the Arabidopsis *SUPPRESSOR OF PHYA-105 1* (*SPA1, AT2G46340*), and for 50% of cotyledon maximum area, *Bo5g001100.1*, an ortholog of *ARABIDOPSIS NAC DOMAIN CONTAINING PROTEIN 2* (*ANAC2, AT1G01720*), were significantly associated with these two traits (**Table 1**). For all these seedling traits, the gene expression levels were negatively correlated for the traits of interest (**Supplementary Figure 3**).

**Figure 3.**
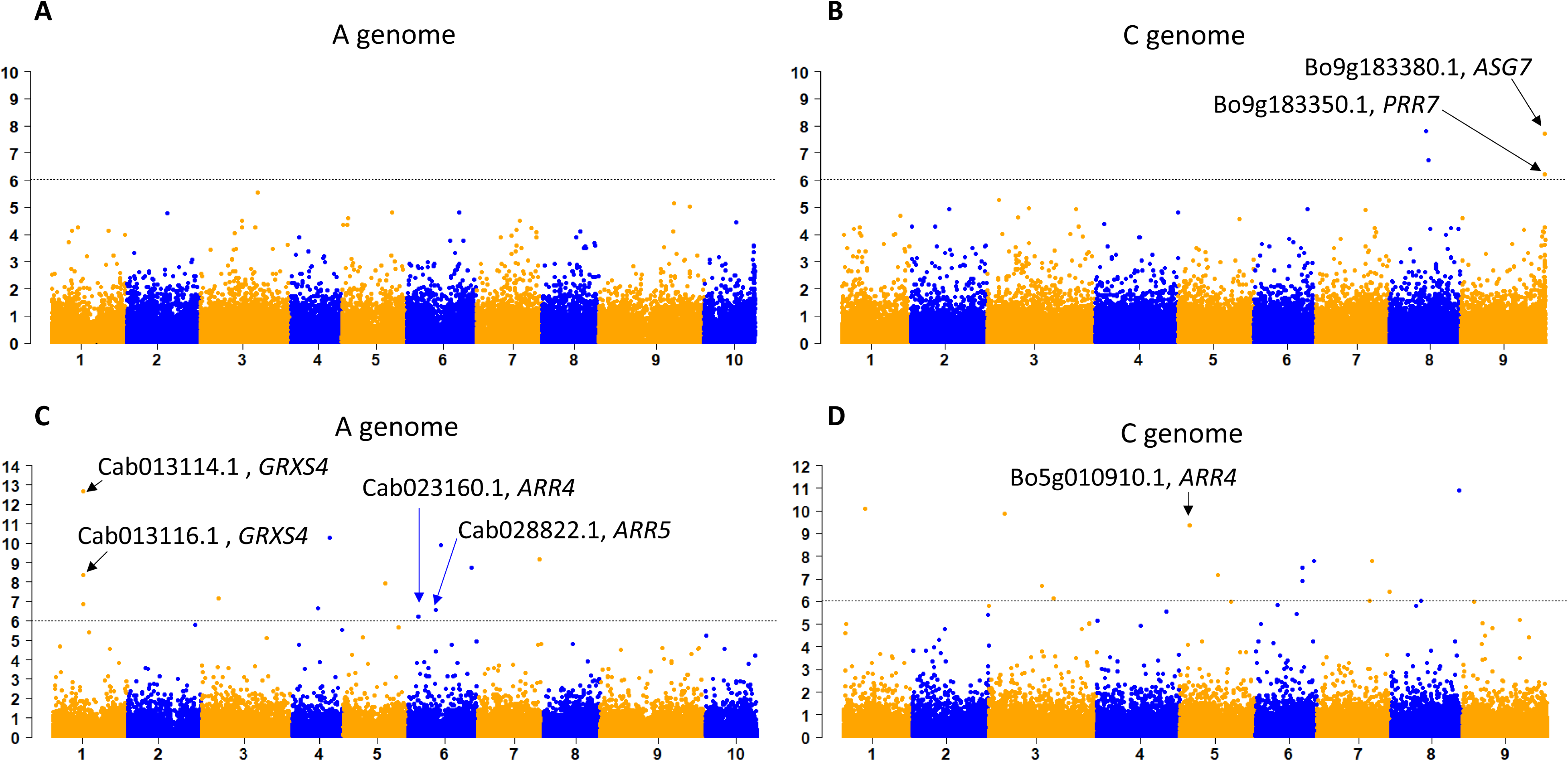
Transcript abundance associated with days to emergence for Chromosome A (A) chromosome C (B) and days to appearance of second leaf for chromosome A (C) and chromosome C (D)genomes of *Brassica napus*. Trait association significance, expressed as -log10P, are plotted on the y-axis. The line marks the Bonferroni threshold.

## Discussion

Seedling vigour has a decisive influence on rapid and uniform seedling emergence and establishment, leading to vigorous crop development and promising final crop yields (Nelson et al., 2022). Here, we studied the phenotypic variation of several seedling traits in an OSR diversity set with an additional eight in-house varieties to understand the relationship of seedling establishment with other seedling development traits. The negative correlation between cotyledon expansion rate and leaf extension for WOSR suggests that the faster the cotyledons expand, the less time the plant needs to reach the leaf extension phase. This strategy is key particularly for WOSR genotypes, as the expansion of cotyledons allow the plants to absorb more sunlight and generate more photoassimilates, leading to increased autumn vigour which is directly related to speed of emergence and fast production of biomass and is critical to the success of the survival of winter oilseed rape crops. The observed phenotypic differences in days to emergence, DAFL and DASL provide insight into the speed of the seedling establishment of different genotypes until the critical stage of two leaves. OSR displays epigeal germination, in which the cotyledons and apical meristem grow through the seedbed to emerge above the surface to form the initial photosynthetic unit (Finch-Savage and Bassel, 2016; Nelson et al., 2022). Therefore, a rapid development of two true leaves is a crucial trait in OSR seedlings as they grow faster than the damage generated by slugs, flea beetles and birds, decreasing its vulnerability to pest attacks. The differences observed in seedling establishment speed were associated with different hypocotyl and root lengths, which are also key traits in OSR. With increasing autumn and spring droughts year after year and climate models predicting an increase of soil moisture deficit for many cropping regions (Vadez et al., 2024) (Kanthavel et al, 2024) (Knox et al., 2010), seedlings require good strategies for water uptake. Most of the low seedling vigour genotypes had short roots, highlighting the importance of the development of the root system for good seedling establishment. On the other hand, the genotypes that most rapidly established the second leaf exhibited almost invariably long hypocotyls, long roots or a combination of long roots and hypocotyls. Long hypocotyls and roots enable deeper sowings and present a higher opportunity of survival (Rebetzke et al., 2007; Thomas et al., 2016). As moisture will become scarcer and more insufficient for seed germination and development near the soil surface, longer roots can reach easily and more efficiently the soil moisture that lies below the topsoil. Within our dataset, BIENVENU DH4 had medium hypocotyl and root lengths compared to the other genotypes from the population. Despite these traits, this genotype still established poorly, as BIENVENU DH4 took the longest time to DASL, showing low seed vigour. This highlights the importance of measuring a combination of seedling traits, as several factors are playing a role in speed of seedling establishment. Although some studies found positive association of seed size and cotyledon area with seed germination rate in OSR (Hatzig et al., 2015), we found weak correlations for seedling components traits measured in this experiment. This is in accordance with Bettey et al. (2000) and Hatzig et al. (2015), who found low correlation of seed size on germination rate and radicle growth on Brassica species.

Determining which genes influence phenotypic traits is a crucial step for crop improvement. The information obtained from the AT analysis identified OSR orthologues of 2 Arabidopsis genes involved in the cytokinin signalling pathway (*ARR4* and *ARR5*) as being significantly associated with seedling establishment speed, specifically DASL. In Arabidopsis, *ARR4* and *ARR5* are A-type ARRs (Arabidopsis Response Regulators) that are activated when they are phosphorylated and negatively regulate cytokinin signalling in negative feedback loop (To et al., 2004; Werner and Schmülling, 2009). *ARR4* and *ARR5* are involved in the control of seed germination by repressing the levels of *ABI5* (Wang et al., 2011; Hatzig et al., 2015). *ARR4* and *ARR5* are also involved in leaf development and expansion. Both genes are negatively regulated by *ANGUSTIFOLIA3 (AN3)*, which stimulates cell division during leaf development in Arabidopsis (Vercruyssen et al., 2014). The repression of these genes by *AN3* could reduce the cytokinin regulated negative inhibition and reinforce this phytohormone signalling, stimulating leaf cell proliferation (Vercruyssen et al., 2014; Vercruysse et al., 2020) (**Figure 4**). *AN3* associates with growth regulator factors (GRF) complexes to regulate the transcription of different target genes. *ARR5* is also repressed by *WUSCHEL* (*WUS*), a transcription factor that is essential for stem cell specification in the shoot apical meristem (Leibfried et al., 2005; Gordon et al., 2009; Ren et al., 2009). *WUS* levels are increased by cytokinin, a phytohormone the biosynthesis of which is in turn promoted by *KNOX* genes. Hence, *ARR5* repression ensures cell division and differentiation in the apical meristem, enhancing the establishment of shoot architecture (Jasinski et al., 2005; Yanai et al., 2005; Souček et al., 2007). The most highly associated GEM with DASL is an ortholog of Arabidopsis *GRXS4*. This gene has been also reported to function downstream of cytokinin signalling, although there are no reports so far for its function in leaves but in root growth instead (Kiba et al., 2005; Patterson et al., 2015). Light and phytochrome play a role in seedling establishment. *SPA1* interacts with *COP1*, and together inhibit and activate transcription factors that promote photomorphogenesis in the dark and in the light, respectively (Chen et al., 2016; Paik et al., 2019), controlling cotyledon expansion. In addition to cytokinins, other hormones also play a role in the speed of seedling establishment. For days to emergence, *ASG7* was significantly associated with the trait., and it has been reported to repress seed germination through ABA and GA signalling (Bassel et al., 2011). *PPR7* was also found associated with days to emergence. This gene interacts with *ABI5* to activate ABA signalling to maintain proper seed germination and post germinative growth. It is also involved with the circadian clock to control seed germination and hypocotyl growth (Li et al., 2020; Yang et al., 2021). Taking also into account that *ARR4* and *SPA1* ortholog are also involved in the control of hypocotyl length in Arabidopsis through phyB and blue light, respectively (Sweere et al., 2001; Mira-Rodado et al., 2007; Rolauffs et al., 2012; Ling et al., 2017; Li et al., 2020), the results suggests a role of hormone and light interplay in the control of seedling establishment.

**Figure 4:**
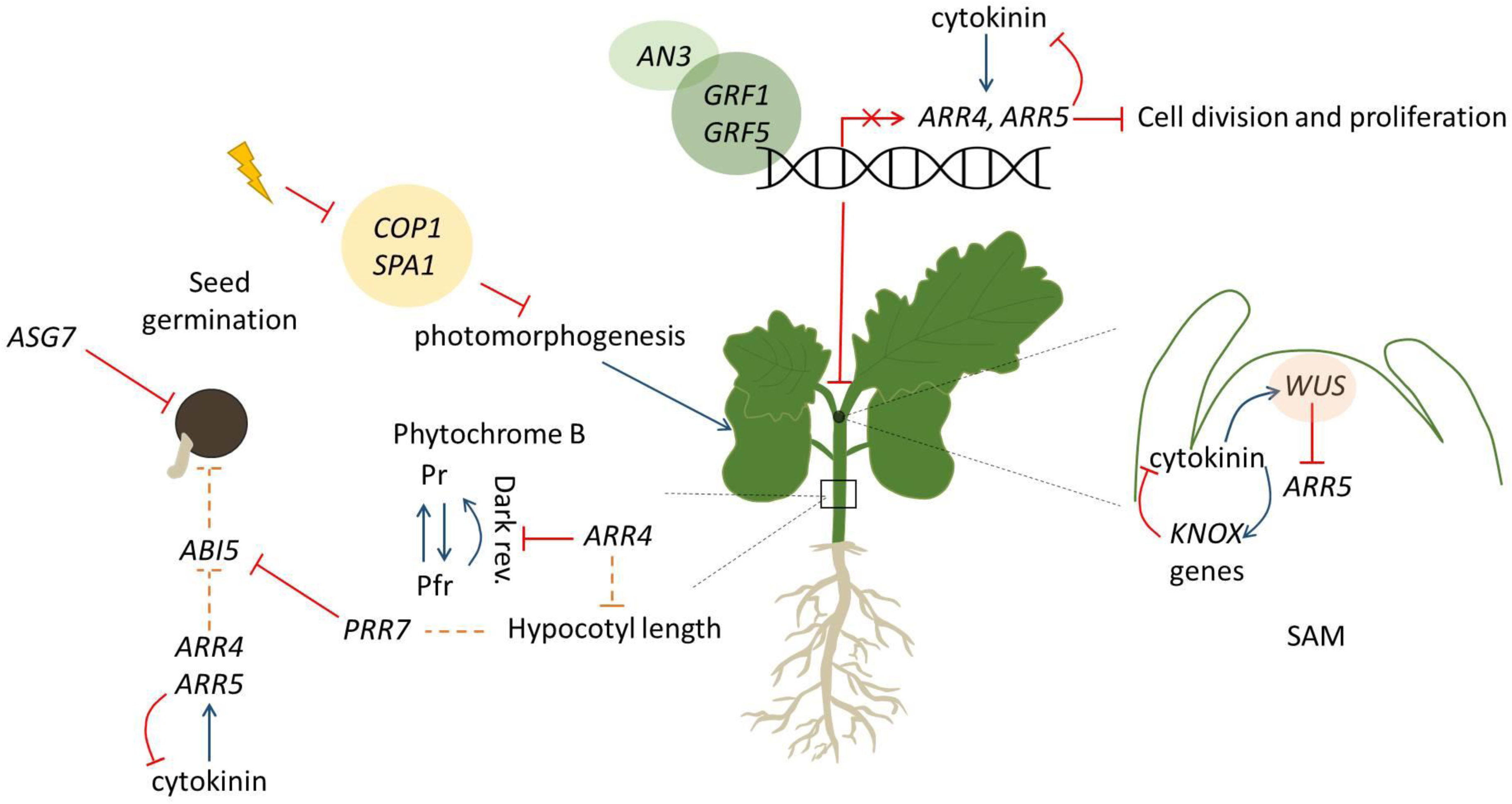
Hypothetical model of seedling establishment for OSR. Function of *ASG7*, *ARR4*, *ARR5* and *PRR7* on seed germination repression. *AN3* and *GFR* complex regulating *ARR4* and *ARR5* transcriptional levels for cell division and proliferation in leaves. *ARR5* levels regulated in the shoot apical meristem (SAM). *WU*S is controlling the levels of stem cells in the SAM (depicted as orange sphere). *COP1* and *SPA1* complex and its role in photomorphogenesis as well as the role of *ARR4* and *PRR7* in hypocotyl length are also depicted for seedling growth. Red lines show inhibition of function. Blue lines depict enhancing of function. Dash orange lines are representing interactions found in literature but not in this study.

While all OSR growers and breeder concur that vigour is an important trait, this is challenging when there is no standardised measurement for vigour. Vigour is not a single measurement in time but a reflection of the growth and development of a WOSR plant seedling over the autumn period. A comprehensive understanding therefore of the underlying traits and their relatedness will allow breeders to gain a better idea of how to measure vigour and subsequently select better OSR varieties.

## Acknowledgements

We thank Andrew Mead (Rothamsted Research, UK) for performing the statistical experimental designs and analysis. We also thank Elsoms seeds for seed for eight pre-breeding lines along with valuable advice and discussions.

