## Supplementary information Table 1 for "Associative transcriptomics in *Brassica napus* suggests a role for Arabidopsis Response Regulator orthologs in seedling vigour"

Supplemental Table S1: List of 95 genotypes included in the diversity set population. The ASSYST code, genotype names, crop type description and the 5 oilseed rape groups are presented.

| ASSYST code | Genotype names | Crop type description | OSR groups |
|---|---|---|---|
| BnASSYST-213 | Abukuma Natane | Winter OSR | Winter OSR |
| BnASSYST-418 | Alta Sweet | Swede | Other |
| BnASSYST-040 | Apex | Modern winter OSR | Winter OSR |
| BnASSYST-090 | Apex-93_5 x Ginyou_3 DH | Winter OSR | Winter OSR |
| BnASSYST-190 | Aphid Resistant Rape | Winter fodder | Other |
| BnASSYST-133 | Baltia | Winter OSR | Winter OSR |
| BnASSYST-091 | Bienvenu DH4 | Winter OSR | Winter OSR |
| BnASSYST-206 | Brauner Schnitikohl | Siberian kale | Other |
| BnASSYST-273 | Bronowski | Spring OSR | Spring OSR |
| BnASSYST-509 | Cabernet | Winter OSR | Winter OSR |
| BnASSYST-510 | Cabriolet | Winter OSR | Winter OSR |
| BnASSYST-185 | Canard | Winter fodder | Other |
| BnASSYST-093 | Canberra x Courage | Winter OSR | Winter OSR |
| BnASSYST-028 | Capitol | Modern winter OSR | Winter OSR |
| BnASSYST-511 | Castille | Winter OSR | Winter OSR |
| BnASSYST-512 | Catana | Winter OSR | Winter OSR |
| BnASSYST-274 | Ceska Krajova | Spring OSR | Spring OSR |
| BnASSYST-207 | Chembere Dzagumhana | Unspecified | Other |
| BnASSYST-513 | Chuanyou 2 | Semiwinter OSR | Semiwinter OSR |
| BnASSYST-137 | Coriander | Winter OSR | Winter OSR |
| BnASSYST-208 | Couve Nabica | Cauve nabica | Other |
| BnASSYST-245 | Cresor | Spring OSR | Spring OSR |
| BnASSYST-261 | Cubs Root | Spring OSR | Spring OSR |
| BnASSYST-514 | Dimension | Winter OSR | Winter OSR |
| BnASSYST-139 | Dippes | Winter OSR | Winter OSR |
| BnASSYST-238 | Drakkar | Spring OSR | Spring OSR |
| BnASSYST-275 | Duplo | Spring OSR | Spring OSR |
| BnASSYST-193 | Dwarf Essex | Winter fodder | Other |
| BnASSYST-194 | English Giant | Winter fodder | Other |
| BnASSYST-263 | Erglu | Spring OSR | Spring OSR |
| BnASSYST-101 | Eurol | Winter OSR | Winter OSR |
| BnASSYST-515 | Excalibur | Winter OSR | Winter OSR |
| BnASSYST-055 | Expert | Modern winter OSR | Winter OSR |
| BnASSYST-516 | Flash | Winter OSR | Winter OSR |
| BnASSYST-218 | Groene Groninger Snijmoes | Unspecified | Other |
| BnASSYST-096 | Hansen x Gaspard DH | Winter OSR | Winter OSR |
| BnASSYST-264 | Helios | Spring OSR | Spring OSR |
| BnASSYST-410 | Huguenot | Swede | Other |
| BnASSYST-517 | Huron x Navajo | Winter OSR | Winter OSR |
| BnASSYST-518 | Inca x Contact | Winter OSR | Winter OSR |
| BnASSYST-107 | Janetzkis Schlesischer | Winter OSR | Winter OSR |
| BnASSYST-411 | Jaune A Collet Vert | Swede | Other |
| BnASSYST-251 | Karat | Spring OSR | Spring OSR |
| BnASSYST-256 | Karoo-057 DH | Spring OSR | Spring OSR |
| BnASSYST-150 | Kromerska | Winter OSR | Winter OSR |
| BnASSYST-108 | Lembkes Malchower (Lenora) | Winter OSR | Winter OSR |

| | | | |
|---|---|---|---|
| BnASSYST-102 | Lesira | Winter OSR | Winter OSR |
| BnASSYST-105 | Licrown x Express DH | Winter OSR | Winter OSR |
| BnASSYST-271 | Liho | Spring fodder | Other |
| BnASSYST-097 | Madrigal x Recital DH | Winter OSR | Winter OSR |
| BnASSYST-160 | Matador | Winter OSR | Winter OSR |
| BnASSYST-268 | Mazowiecki | Spring OSR | Spring OSR |
| BnASSYST-186 | Moana, Moana Rape | Winter fodder | Other |
| BnASSYST-257 | Monty-028 DH | Spring OSR | Spring OSR |
| BnASSYST-259 | N02D-1952 | Spring OSR | Spring OSR |
| BnASSYST-520 | Ningyou 7 | Semiwinter OSR | Semiwinter OSR |
| BnASSYST-258 | N01D-1330 | Spring OSR | Spring OSR |
| BnASSYST-109 | Norin | Winter OSR | Winter OSR |
| BnASSYST-521 | Palmedor | Winter OSR | Winter OSR |
| BnASSYST-522 | POH 285, Bolko | Winter OSR | Winter OSR |
| BnASSYST-204 | Q100 | Synthetic | Other |
| BnASSYST-523 | Quinta | Winter OSR | Winter OSR |
| BnASSYST-098 | Rafal DH1 | Winter OSR | Winter OSR |
| BnASSYST-209 | Ragged Jack | Rape kale | Other |
| BnASSYST-168 | Ramses | Winter OSR | Winter OSR |
| BnASSYST-221 | Rapid Cycling Rape (CrGC5) | Rape kale | Other |
| BnASSYST-524 | Rocket | Winter OSR | Winter OSR |
| BnASSYST-113 | Samourai | Winter OSR | Winter OSR |
| BnASSYST-414 | Sensation Nz | Swede | Other |
| BnASSYST-106 | Shannon x Winner DH | Winter OSR | Winter OSR |
| BnASSYST-526 | Shengliyoucai | Semiwinter OSR | Semiwinter OSR |
| BnASSYST-211 | Siberische Boerenkool | Siberian kale | Other |
| BnASSYST-212 | Slapska, Slapy | Unspecified | Other |
| BnASSYST-172 | Slovenska Krajova | Winter OSR | Winter OSR |
| BnASSYST-239 | Stellar DH | Spring OSR | Spring OSR |
| BnASSYST-260 | Surpass400-024 DH | Spring OSR | Spring OSR |
| BnASSYST-229 | SWU Chinese 1 | Semiwinter OSR | Semiwinter OSR |
| BnASSYST-230 | SWU Chinese 2 | Semiwinter OSR | Semiwinter OSR |
| BnASSYST-203 | Taisetsu | Winter vegetable | Other |
| BnASSYST-269 | Tantal | Spring OSR | Spring OSR |
| BnASSYST-099 | Tapidor DH | Winter OSR | Winter OSR |
| BnASSYST-527 | Temple | Winter OSR | Winter OSR |
| BnASSYST-436 | Tina | Swede | Other |
| BnASSYST-283 | Topas | Spring OSR | Spring OSR |
| BnASSYST-307 | Tribune | Spring OSR | Spring OSR |
| BnASSYST-053 | Verona | Modern winter OSR | Winter OSR |
| BnASSYST-401 | Vige DH1 | Swede | Other |
| BnASSYST-528 | Vision | Winter OSR | Winter OSR |
| BnASSYST-270 | Weihenstephaner | Spring OSR | Spring OSR |
| BnASSYST-240 | Westar DH | Spring OSR | Spring OSR |
| BnASSYST-394 | Willi | Spring OSR | Spring OSR |
| BnASSYST-529 | Xiangyou 15 | Semiwinter OSR | Semiwinter OSR |
| BnASSYST-438 | York | Swede | Other |
| BnASSYST-530 | Zhongshuang II | Semiwinter OSR | Semiwinter OSR |
| BnASSYST-237 | Zhouyou | Semiwinter OSR | Semiwinter OSR |
