## Supplementary Figure 1 for "Associative transcriptomics in *Brassica napus* suggests a role for Arabidopsis Response Regulator orthologs in seedling vigour"

**Supplementary Figure 1.** A) Plates disposition inside a SANYO cabinet. Plates sealed in light tight black bags in batches of 32 arranged upright are on the left-hand side . Plates in statistical randomised disposition and exposed to light conditions on right-hand side. B) Germination plate after 3 days in the dark. A total of 20 seeds per genotype were placed in each row, with a total of 15 rows in total per plate.

A B

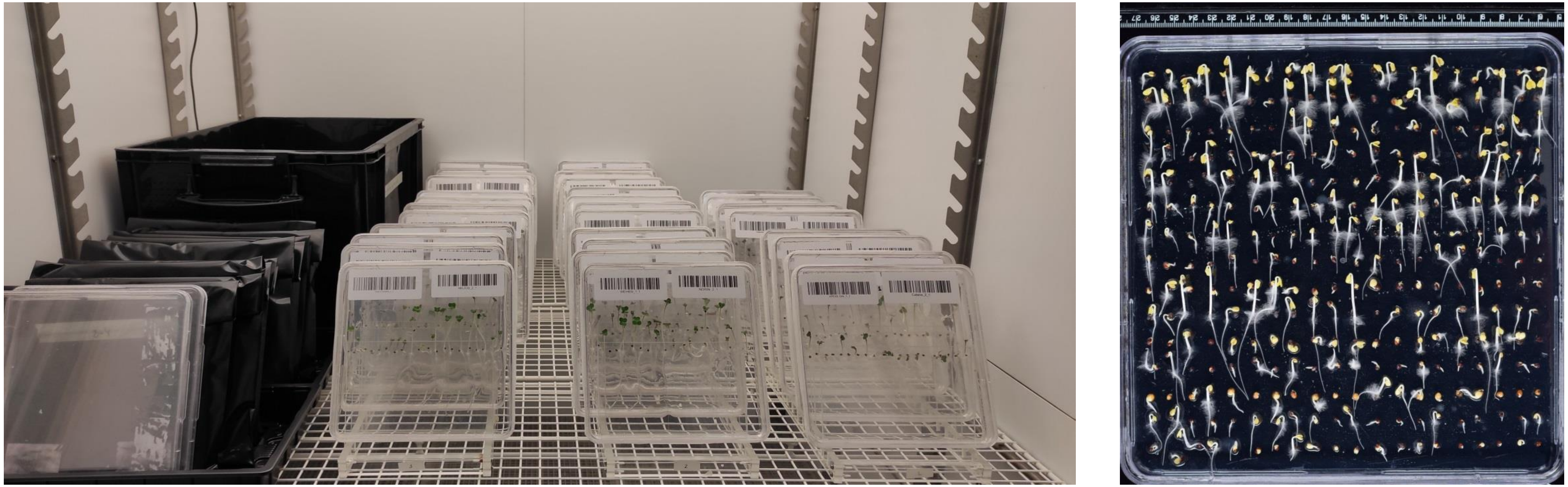

**Supplementary Figure 2.** Pearson's correlations for A) 103 genotypes studied in the population, B) 42 Winter OSR genotypes and C) 22 Spring OSR genotypes. DAFL: Days to appearance of first leaf. DASL: Days to appearance of second leaf. * Indicates statistically significant correlations with values higher than 0.50.

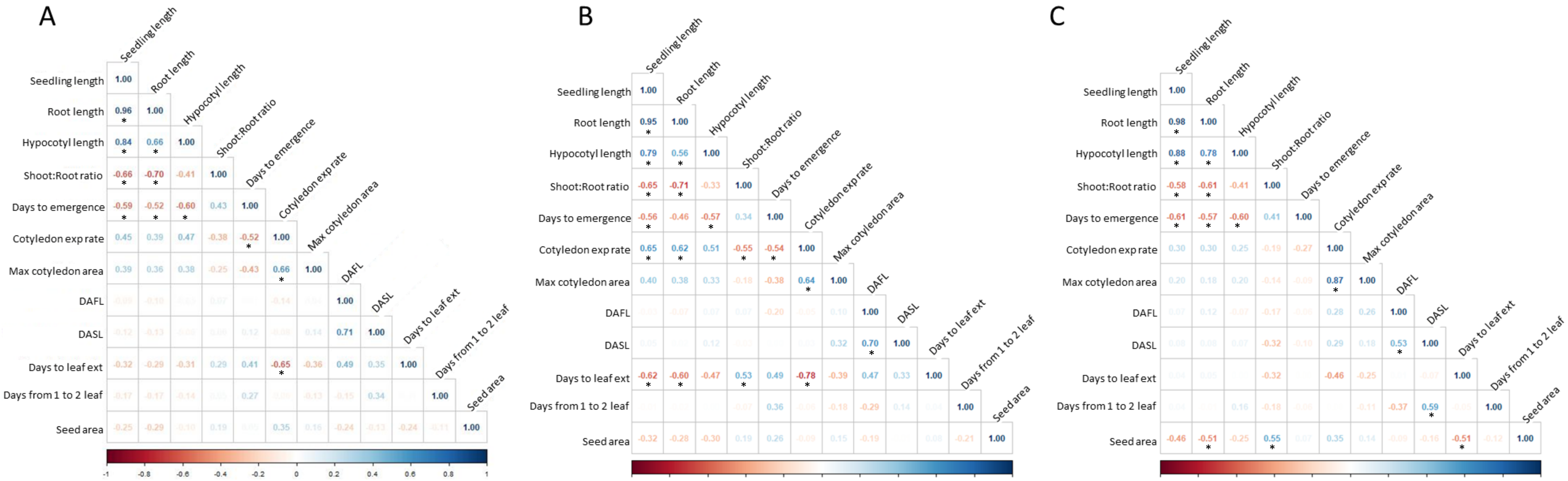
