## Supplementary Figure 3 for "Associative transcriptomics in *Brassica napus* suggests a role for Arabidopsis Response Regulator orthologs in seedling vigour"

**Supplementary Figure 3.** Gene expression levels of significant orthologs associated with days to emergence for *ASG7* (A) and *PRR7* (B), gene expression for *ARR4* (C-D), *ARR5* (E), *GRXS4* (F-G), *SPA1* (H) and *ANAC2* (I) for days to reach 25% or 50% of cotyledon maximum area respectively. Plots were generated in R using ggplot2 package based on a liner model of the adjusted means versus RPKM (Reads Per Kilobase per Million mapped reads). Correlation coefficient and *P*-values are shown in each graph .

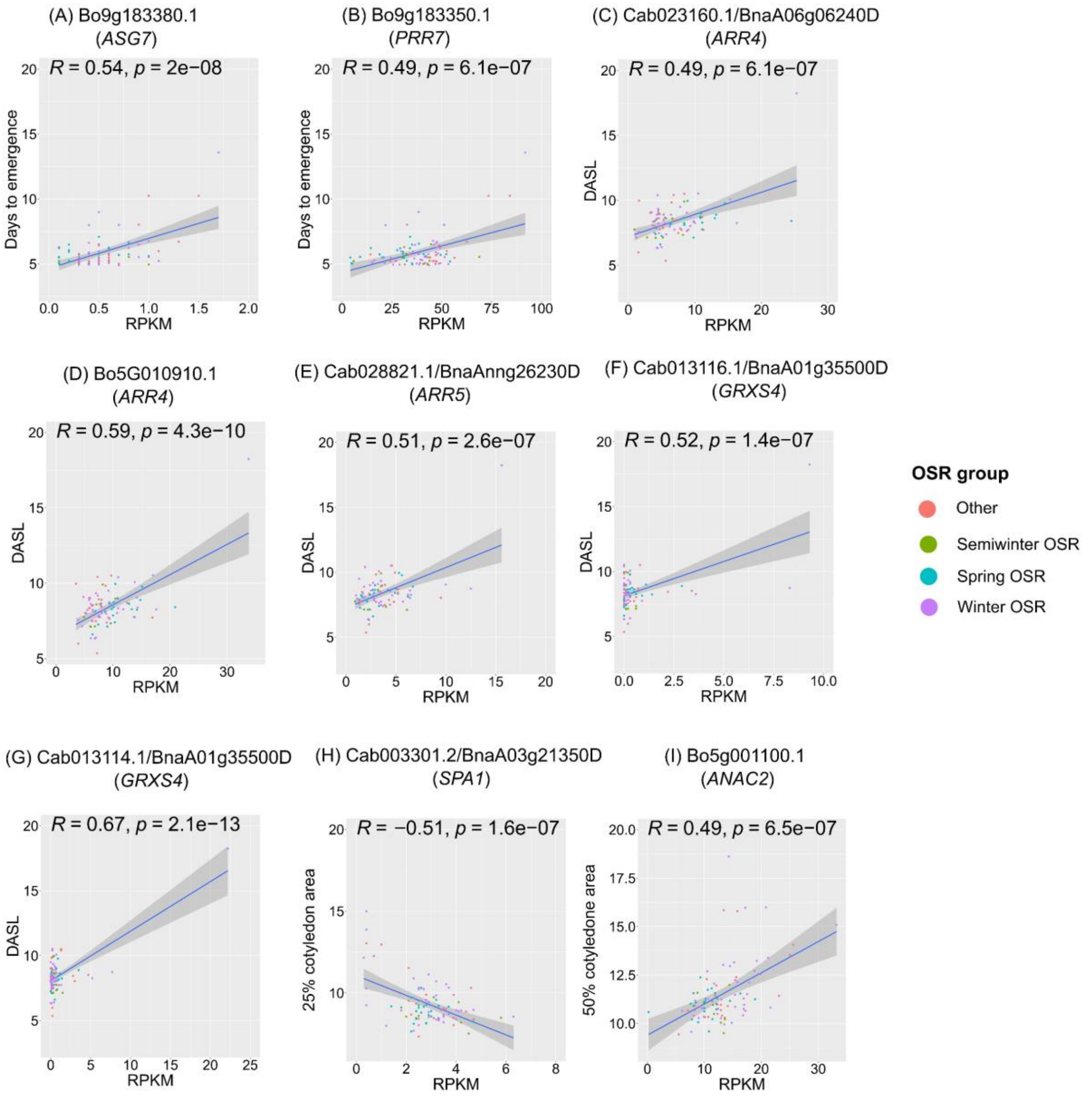
